# Expanded Distribution and Predicted Suitable Habitat for the Critically Endangered Yellow-tailed Woolly Monkey (*Lagothrix flavicauda*) in Peru

**DOI:** 10.1101/2022.05.19.492669

**Authors:** Melissa A. Zarate, Sam Shanee, Elvis Charpentier, Yeissy Sarmiento, Christopher A. Schmitt

## Abstract

The Tropical Andes Biodiversity Hotspot holds a remarkable number of species at risk of extinction due to anthropogenic habitat loss, hunting and climate change. One of these species, the Critically Endangered yellow-tailed woolly monkey (*Lagothrix flavicauda*), was recently sighted in Junín region, 206 kilometres south of its previously known distribution. The range extension, combined with continued habitat loss, calls for a re-evaluation of the species’ distribution and available suitable habitat. Here, we present novel data from surveys at 53 sites in the regions of Junín, Cerro de Pasco, Ayacucho and Cusco. We encountered *L. flavicauda* at 9 sites, all in Junín, and the congeneric *L. l. tschudii* at 20 sites, but never in sympatry. Using these new localities along with all previous geographic localities for the species, we made predictive Species Distribution Models based on Ecological Niche Modelling using a generalized linear model and maximum entropy. Each model incorporated bioclimatic variables, forest cover, vegetation measurements, and elevation as predictor variables. Model evaluation showed >80% accuracy for all measures. Precipitation was the strongest predictor of species presence. Habitat suitability maps illustrate potential corridors for gene flow between the southern and northern populations, although much of this area is inhabited by *L. l. tschudii*. An analysis of the current protected area (PA) network showed ∼47% of remaining suitable habitat is unprotected. With this, we suggest priority areas for new protected areas or expansions to existing reserves that would conserve potential corridors between *L. flavicauda* populations. Further surveys and characterization of the distribution in intermediate areas, combined with studies on genetic flow, are still needed to protect this species.

## Introduction

Changes in species’ distributions are of major concern for conservationists due to the continual expansion of human populations and land use (Hansen *et al*. 2001; Newbold *et al*. 2014; Dai *et al*. 2021) that shift, contract, or fragment a species’ geographic range. Such altered ranges may isolate populations, enhancing the likelihood of inbreeding depression or local extinction (Ewers and Didham 2005; Calkins *et al*. 2021; Solórzano-Garcia *et al*. 2021). Information on the relative distributions of different populations of a species can reveal what types of landscapes or species community composition—in the case of competitive exclusion—act as geographic or ecological barriers to gene flow (Case and Taper 2000; Blair and Melnick 2012; Sales *et al*. 2019; Pázstor *et al*. 2020). To conserve natural biodiversity in the face of expanding land use by humans, conservation management must operate on a landscape scale, understanding species-ecosystem relationships and species’ responses to habitat change (Bellamy *et al*. 2013; Robillard *et al*. 2015; Xioa *et al*. 2019). Informed management planning involves understanding the true current distribution of a species and the necessary habitat components to predict how distribution changes will occur in the future, and the best possible options for area protection.

The yellow-tailed woolly monkey (*Lagothrix flavicauda*) is considered Critically Endangered and amongst the most threatened primate species in the world (Mittermeier *et al*. 2012; Shanee *et al*. 2021), making it a key focus of conservation initiatives in northern Peru (Shanee *et al*. 2018). The Tropical Andes Biodiversity Hotspot (TABH), to which the species is endemic, is among the top five hotspots predicted to lose the most biological diversity due to continued anthropogenic activities (Brooks *et al*. 2002). In northern Peru, the greatest threat to species and habitats in the TABH is habitat destruction for cattle ranching, logging, and other, mainly small scale, economic activities, fuelled by a growing human population and aided by the construction of highways and incentivized access to lands (Oliveira *et al*. 2007; Shanee 2011; Shanee 2012; Programa Bosques 2015; GIZ 2016; Laurance 2018; Shanee and Shanee 2016). With this migrant economic development, roads are continuously built to access new areas, creating a cycle of increasing deforestation for the extraction of resources and settlement (Gallice, Larrea-Gallegos and Vázquez-Rowe 2019). Hunting of primates for bushmeat and the pet trade is also putting *L. flavicauda* and other species at further risk (Shanee 2011; Shanee 2012; Shanee *et al*. 2017).

The geographic range of *L. flavicauda* – long thought to be restricted to the northern regions of Amazonas and San Martín and neighbouring areas of Huánuco, La Libertad, and Loreto – has been continuously re-evaluated through survey efforts (Mittermeier, de Macedo-Ruiz, Luscombe 1975; Graves *et al*. 1980; Leo Luna 1980; Parker and Barkley 1981; Butchart *et al*. 1995; Shanee *et al*. 2007; Shanee 2016; Shanee 2011). Recent observations in the region of Huánuco have found *L. flavicauda* populations as far as the south eastern border with the region of Pasco (Aquino *et al*. 2015; Aquino, Garcia and Charpentier 2016). Camera trap observations led to the discovery of a new population of *L. flavicauda* in 2018 (McHugh *et al*. 2020). This new population, in the Inchatoshi Kametsha Conservation Concession, Junín, is about 206 km south of the previously known southern range limit for the species (McHugh *et al*. 2020), with no sightings in intermediate areas.

*Lagothrix flavicauda* is generally restricted to montane forests between 1,400 to 2,800 m above sea level (m.a.s.l.) along the eastern slopes of the Andes (Aquino *et al*. 2017, Shanee 2011), with occasional occurrences below this elevation (Allgas *et al*. 2014; Paterson and Lopez Wong 2014). Their preferred habitat depends on local climate and forest composition (Shanee 2016; Almeyda-Zambrano *et al*. 2019), and they are able to survive at least for a time in moderately disturbed habitats when hunting pressure is low (Aquino *et al*. 2015; Shanee and Shanee 2015). This may be, in part, due to a flexible diet consisting of flowers, leaf petioles, epiphytic roots, vertebrates, and soil, although they are predominantly frugivorous. Fruits make up almost half of their dietary intake, which is dominated by a handful of species and genera with large fleshy fruits (Shanee and Shanee 2011, Shanee 2014; Fack *et al*. 2018), making food tree density and phenology important factors for sustaining viable populations. Previous studies suggest that the protected area (PA) network in northern Peru is insufficient to protect *L. flavicauda* from the consequences of human population growth, habitat degradation, and climate change (Buckingham and Shanee 2009; Shanee 2016). As the number of PAs in Peru change, along with our knowledge of primate species distributions, it is critical to re-assess their coverage with current data.

Many conservation and research initiatives use *L. flavicauda* as a flagship species in Peru (Shanee *et al*. 2018). Although these have led to an increasing number of private, communal, and state PAs within the species distribution (Shanee *et al*. 2017; Shanee *et al*. 2020), they have predominantly focused on populations in Amazonas and San Martín (Shanee and Shanee 2015). Field surveys are vital to determining the species’ distribution and habitat preferences outside these well-studied regions; however, they are often expensive, time consuming, and can be unfeasible depending on the terrain, access, and public order limitations (Young 1996; Shanee and Shanee 2016). Predictive species distribution modelling (SDM) can highlight optimal areas for field surveys through comparison of the ecological conditions found within a given species’ known range, with those across a wider area, showing where conditions are most favourable for the species presence (Phillips, Anderson and Schapire 2006; Ramirez-Villegas *et al*. 2014; Guisan, Thuiller and Zimmerman, 2017).

Species’ distributions are predicated on three main conditions: the species must have the means to disperse into and out of a habitat, the habitat must have the correct combination of environmental variables to make it suitable, and the abiotic conditions must be able to maintain necessary species interactions (Guisan *et al*. 2017). SDM considers the second condition by using known habitat parameters to determine which are most important to the distribution of the focal species, and uses them to locate potentially suitable habitat (Dong *et al*. 2019; Cianfranni *et al*. 2010; Liu *et al*. 2019). Such modeling may be used to prioritize search efforts when financial or human resources are not available for surveying a region of interest in person. For endangered species, analyzing correlations between these variables and known species occurrences can highlight conditions which the species requires for its long-term survival in the region of interest, and can be used to develop more effective conservation strategies. With this, conservationists can determine why a species may not be present in a particular area (Liu *et al*. 2019) or increase research and conservation effort in areas that appear to be suitable (LaRue and Nielsen 2008).

Here we compare the results of two SDM approaches—Generalized Linear Modelling (GLM) and Maximum Entropy (MaxEnt)—using the results of our recent field surveys in central and southern Peru. We created SDMs combining these novel localities with all previous records to create the most complete SDM for the species to date. We use the results of these approaches to highlight remaining suitable *L. flavicauda* habitat, particularly in its less-studied southern range. Finally, we evaluate the current Peruvian PA network’s coverage of suitable habitat within the species’ distribution, noting priority areas for new PAs, conservation corridors, and future surveys between the northern and recently-discovered Junín populations.

## Methods

### Population Surveys

We conducted five surveys between May 2019 and May 2021. Due to Covid-19 related travel restriction, no surveys were conducted between May and December 2020, with the May 2020 survey being cut short due to implementation of national quarantine measures. Surveys were carried out in the regions of Ayacucho, Cerro de Pasco, Cusco, and Junín. Survey sites were selected based on preliminary MaxEnt models which used previously published localities, and updated with new sightings as the study progressed, with sites selected considering access routes, land ownership and researcher safety.

Survey efforts followed methods of previous surveys for the species (Shanee 2011). We gathered locality data along existing trail systems with local residents as field guides. Occasionally new trails were opened to enter new areas; however, this was typically avoided to minimize habitat disturbance. We visited sites for 2-9 days, with effort determined by available habitat size, and the possibility of *L. flavicauda* being found or confirmation of its presence. We recorded localities and points of visual or audio detection of all primates with a handheld GPS. We also gathered secondary evidence from local informants in and around the areas visited. Using images and verbal descriptions of *Lagothrix* spp. we cross-referenced information from multiple informants at each site.

### Species Distribution Modelling

We selected two standard SDM methods— GLM and MaxEnt. To avoid overestimation of suitable habitat, we limited prediction outputs to between -80° W and -65° W, and -5° S and -15° S. We filtered this to only include areas between 1,000 and 3,500 m.a.s.l., i.e., within the species’ altitudinal range, which includes the entirety of the eastern slopes of the Peruvian Andes. Models used presence points from our own surveys combined with localities from recent published studies, i.e., ≤10 years (Shanee 2011; Allgas *et al*. 2015; Aquino *et al*. 2016; Aquino *et al*. 2017; McHugh *et al*. 2020), and an additional six localities from the Global Biodiversity Information Facility (GBIF Secretariat 2021) (Fig. 1; Table S1).

**Fig. 1.**
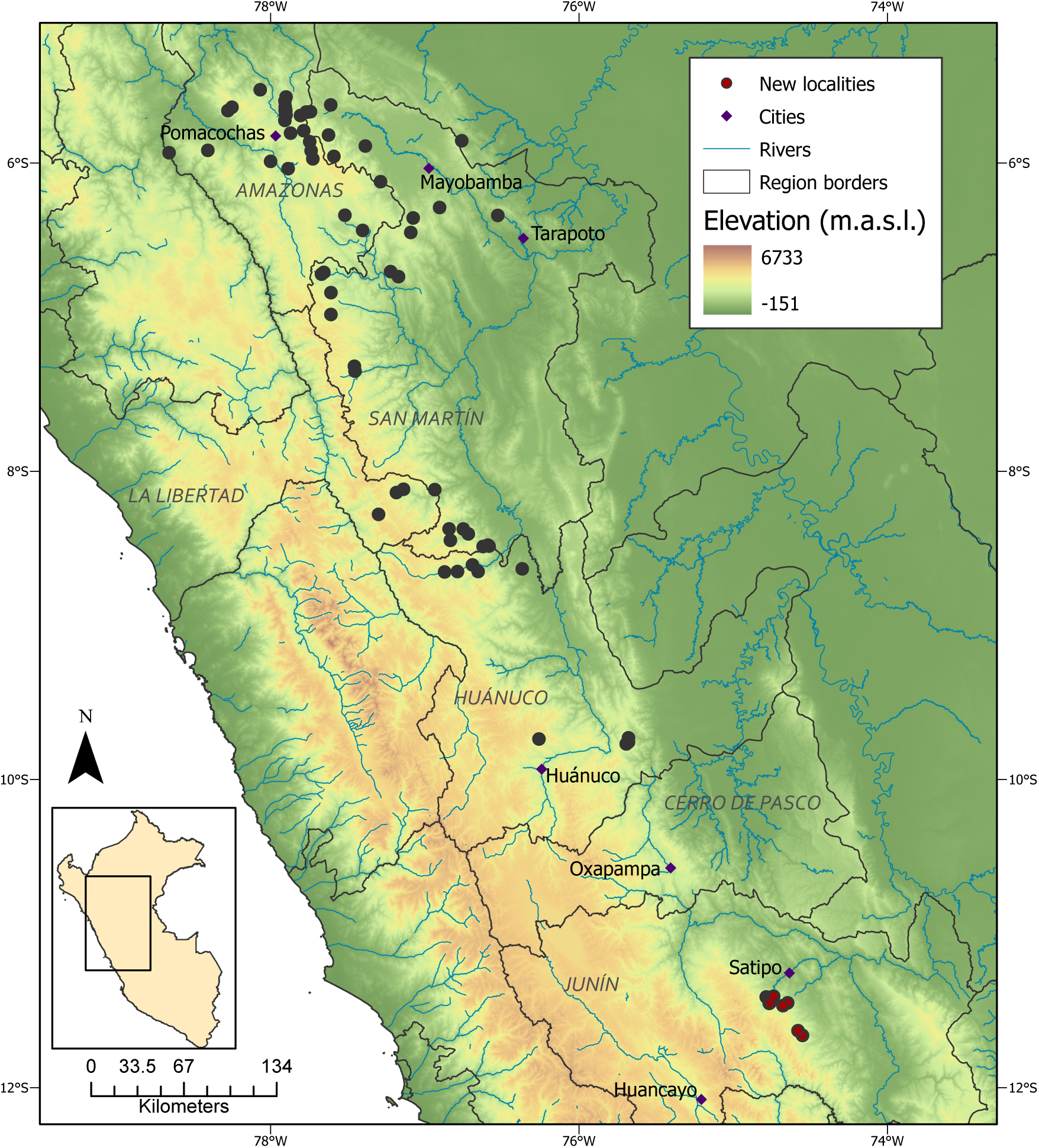
*Lagothrix flavicauda* occurrences used as presence points in habitat suitability modelling (HSM) in this study.

Environmental data were all from publicly available sources (Table S2). We used the 19 bioclimatic variables from *WorldClim2*, available in the R package raster v. 3.1-5 (Hijmans *et al*. 2020) and elevation data from the NASA Shuttle Radar Topography Mission (SRTM; v. 4) at 30-meter resolution. We used forest cover data from the Global Forest Change (v. 1.7) dataset of percent tree cover from the year 2000, in which tree cover is given as a gradient of percentage canopy connectivity per 30 m pixel for vegetation taller than 5 m. Forest loss and gain between the years 2000 and 2018 were also analyzed, but were excluded from the model due to an insufficient number of areas experiencing any recorded loss or gain (fewer than 10 pixels within the study region, which did not coincide with the location of any presence points). We also used the Vegetation Health Product (VHP) dataset, at 1 km resolution, from the NOAA Centre for Satellite Applications and Research, which has been created from an algorithm providing estimates of moisture, thermal, and wind conditions in combination with infrared imaging to achieve an estimate of vegetation conditions in response to weather impacts. All data layers were re-sampled to 30 m resolution.

### Generalized Linear Model

All GLMs were evaluated in R (v. 3.5.2; R Core Team 2018). We modelled the ecological niche of *L. flavicauda* as a binomial response variable of presence/pseudoabsence using a binomial variance and logistic link function. We randomly selected pseudoabsence points to counter sampling bias from the lack of absence data existing for the species using the *randomPoints* function in the *dismo* package v.1.1-4 (Hijmans *et al*. 2017), generating 355 in the study area within an elevation mask including only areas within the altitudinal range. All presence points were excluded from the background point extraction process. We then extracted and standardized values from all environmental layers at each pseudoabsence and presence point to be used in the models. Models were built excluding correlated predictor variables based on Pearson’s *r* value (± 0.75). Our final model was selected using the corrected Akaike Information Criterion (AICc) procedure (Akaike 1973; Hurvich and Tsai 1993).

We evaluated the predictive ability of our model using the area under the ROC curve (AUC), which summarizes model accuracy by giving the probability that the model ranks random presence sites over pseudoabsence sites. We calculated the AUC by building the models from a training set, and then applying it to the remainder of the data as a test set to evaluate predictive performance (Fielding and Bell 1997; Hirzel *et al*. 2006). Due to the relatively small number of presence points, we bootstrapped this process 1000 times using multiple distributions of presence data in the training and testing sets (Hein *et al*. 2007; Guisan *et al*. 2017). We also tested model accuracy using Cohen’s kappa statistic and percent accuracy from a confusion matrix as a threshold-dependent method of evaluation (Guisan *et al*. 2017). Using training and test data, we set a probability threshold so that any point with a presence probability of 0.4 or higher was marked as a presence when establishing a confusion matrix of model predictions. The kappa statistic value ranges from -1 (complete disagreement between predicted and actual values) to 0 (predictions equated to random chance), to +1 (complete agreement with actual values).

### Maximum Entropy Model

The MaxEnt model was created and evaluated in MaxEnt Programming Software (v. 3.4.4; Phillips 2006) and re-evaluated in R. We applied default settings to our MaxEnt run, with the following exceptions: background predictions were written for evaluation and mapping purposes and we selected a random test percentage of 25% of our presence points. We used the same non-correlated variables as in the GLM. MaxEnt measures variable contribution by permutation importance determined by randomly permuting the values of each variable among the training points, measuring the resulting decrease in training AUC, and normalizing these values to give a percentage (Phillips *et al*. 2006). MaxEnt evaluates the model using a ROC curve and calculating the model AUC. We corroborated the given value by bootstrapping the AUC calculation 1000 times in R using the background prediction values.

### Suitable Habitat Classification and Protected Area Assessment

To assess habitat suitability, we assigned habitat based on the predicted probability of species presence to categories of “Good” (P(robability of species presence) between 0.25 and 0.75), “Very Good” (P>0.75), and “Low” (P<0.25), with “Good” and “Very Good” being considered suitable habitat. To incorporate the probable impact of hunting into our results, we considered habitat within 1 km of human settlement to have high hunting pressure for *L. flavicauda* (Shanee 2016). We used data from Humanitarian OpenStreetMap Team (2020) to distinguish human settlements, and deemed any suitable habitat within a 1 km buffer of settlement as low suitability. We overlaid the suitable habitat predicted by both models to find where the predictions intersected. Finally, we conducted an analysis of the PA coverage of suitable *L. flavicauda* habitat by overlaying the Peruvian PA network from the World Database of Protected Areas (WDPA; UNEP-WCMC 2020) with the habitat predictions and calculated the percent of habitat that was considered Good or Very Good within the network. This was repeated with the network of non-governmental conservation concessions in Peru, downloaded from the National Forestry Service (SERFOR), as these concessions are not included in the WDPA files.

### Data Accessibility

All open-source data used for modelling are openly accessible via the links provided in the references. Presence data, prediction results, and all code can be found at https://doi.org/10.5061/dryad.dz08kps0g.

## Results

### Population Surveys

In total we surveyed 53 new sites in central and southern Peru. We encountered *L. flavicauda* at 9 of these sites, all to the north of the Mantaro river in Junín (Fig. 2). In surveys south of this point we did not find any evidence of the species, from our own surveys or from local informants, suggesting a possible southern limit for the species between ∼11.6 and ∼12° south (Table 1). We recorded the congeneric Peruvian woolly monkey (*L. lagotricha tschudii*) at 20 sites, both north and south of the southern *L. flavicauda* population. At no sites were the species sympatric.

**Fig. 2.**
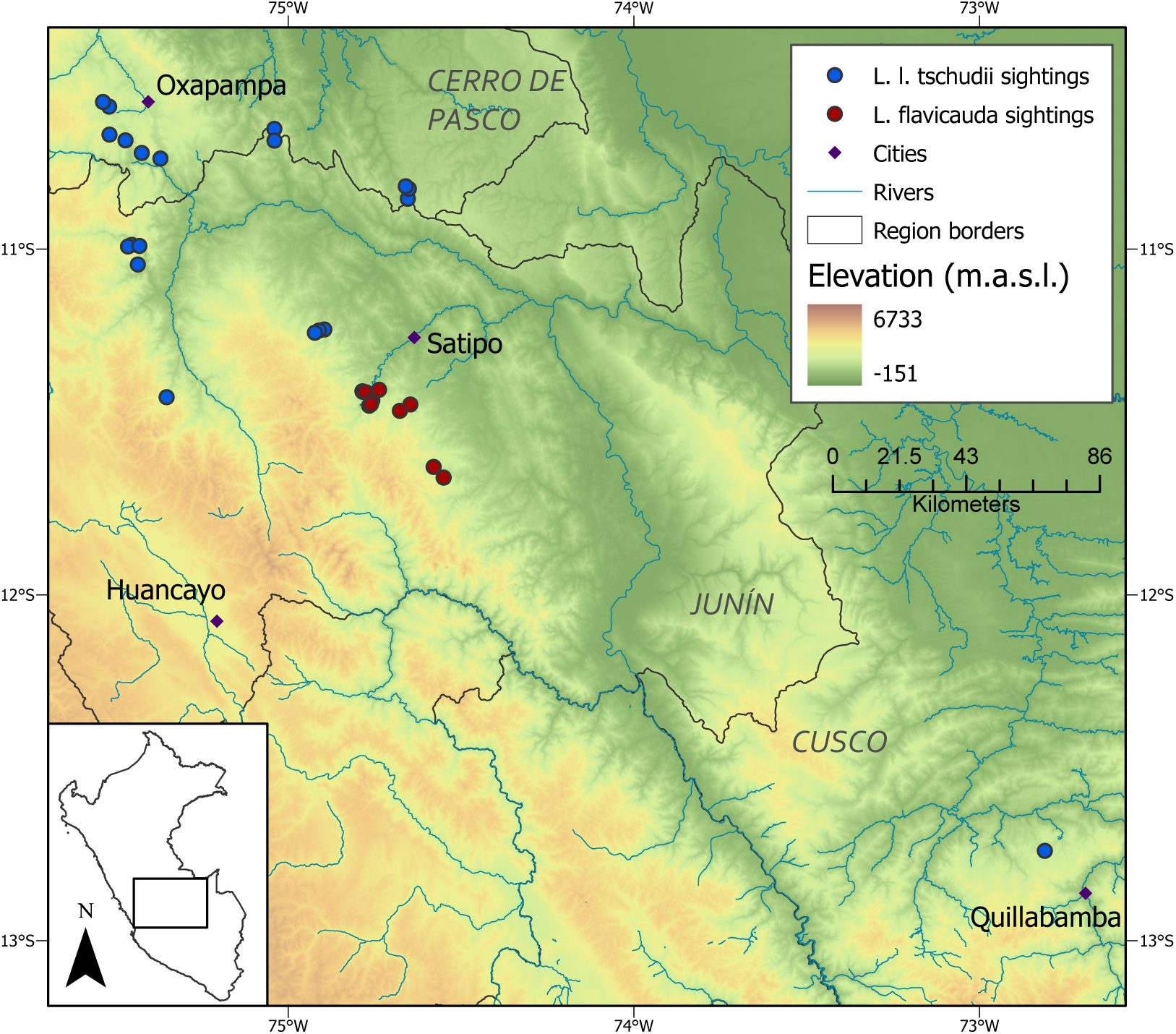
Survey results for *L. flavicauda* and *L. lagotricha tschudii*.

**Table 1.** Coordinates where *L. flavicauda* and *L. l. tschudii* were found in our surveys.

| Region | Species | Longitude | Latitude | m.a.s.l. |
| --- | --- | --- | --- | --- |
| Junín | *L. flavicauda* | -74.7659 | -11.4530 | 2179 |
| Junín | *L. flavicauda* | -74.7568 | -11.4378 | 1869 |
| Junín | *L. flavicauda* | -74.7609 | -11.4490 | 1866 |
| Junín | *L. flavicauda* | -74.6467 | -11.4496 | 1609 |
| Junín | *L. flavicauda* | -74.6770 | -11.4678 | 1825 |
| Junín | *L. flavicauda* | -74.7609 | -11.4490 | 1866 |
| Junín | *L. flavicauda* | -74.5504 | -11.6610 | 2012 |
| Junín | *L. flavicauda* | -74.5793 | -11.6301 | 2067 |
| Junín | *L. flavicauda* | -74.7372 | -11.4069 | 1422 |
| Pasco | *L. l. tschudii* | -75.4709 | -10.6859 | 1410 |
| Pasco | *L. l. tschudii* | -75.3698 | -10.7384 | 1257 |
| Pasco | *L. l. tschudii* | -75.4238 | -10.7223 | 1323 |
| Pasco | *L. l. tschudii* | -75.5182 | -10.5882 | 1612 |
| Pasco | *L. l. tschudii* | -75.5358 | -10.5748 | 2088 |
| Pasco | *L. l. tschudii* | -75.5171 | -10.6693 | 2349 |
| Junín | *L. l. tschudii* | -75.4529 | -10.9879 | 1771 |
| Junín | *L. l. tschudii* | -75.4635 | -10.9928 | 2490 |
| Junín | *L. l. tschudii* | -75.4307 | -10.9911 | 1622 |
| Junín | *L. l. tschudii* | -75.4352 | -11.0447 | 1672 |
| Junín | *L. l. tschudii* | -75.3518 | -11.4285 | 2365 |
| Junín | *L. l. tschudii* | -74.8955 | -11.2327 | 1609 |
| Junín | *L. l. tschudii* | -74.9112 | -11.2358 | 1795 |
| Junín | *L. l. tschudii* | -74.9229 | -11.2421 | 1080 |
| Pasco | *L. l. tschudii* | -74.6540 | -10.8546 | 1465 |
| Pasco | *L. l. tschudii* | -74.6510 | -10.8259 | 1410 |
| Pasco | *L. l. tschudii* | -74.6600 | -10.8186 | 1257 |
| Pasco | *L. l. tschudii* | -75.0401 | -10.6526 | 1323 |
| Pasco | *L. l. tschudii* | -75.0399 | -10.6866 | 1612 |
| Cusco | *L. l. tschudii* | -72.8110 | -12.7401 | 1948 |

### Species Distribution Modelling

After the multicollinearity reduction, the GLM with the lowest AICc (242) contained elevation, mean diurnal temperature range, isothermality, precipitation of wettest month, precipitation seasonality, precipitation of coldest quarter, percent forest cover and VHP as predictor variables. These were the variables used in both model types. The suitable habitat predicted by the two models overlapped by 47.5%, with the GLM being more conservative in the southern regions (Fig. 3).

**Fig 3.**
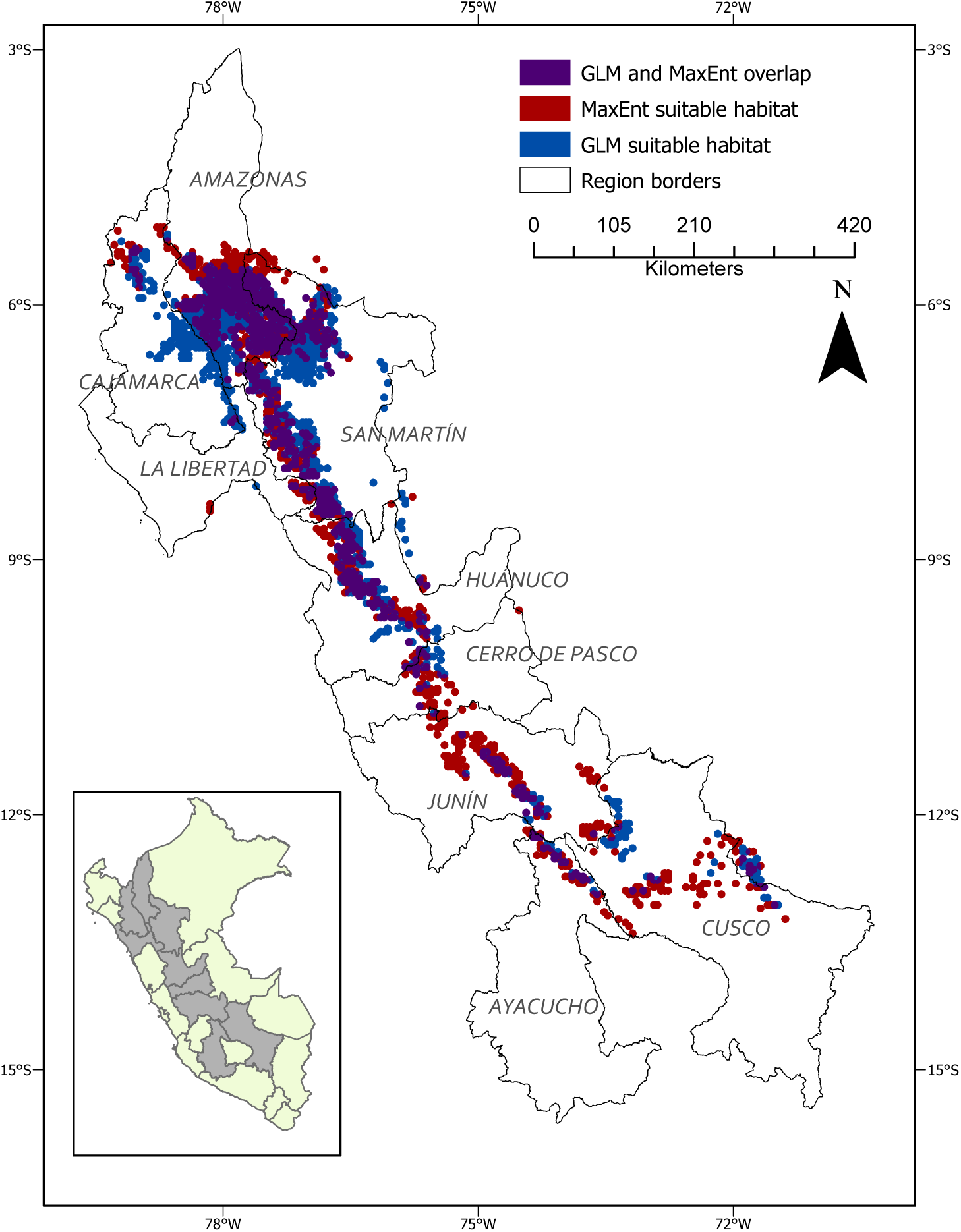
Suitable habitat (Good and Very Good) as indicated by the GLM, MaxEnt model, and their overlap.

### Generalized Linear Modelling

The GLM found all predictor variables to be significantly associated with the likelihood of *L. flavicauda* presence with the exceptions of isothermality and forest cover. The variable most highly associated with *L. flavicauda* habitat was precipitation seasonality, which had a negative relationship. Precipitation of the wettest month had the strongest positive correlation (Table 2).

**Table 2.** Correlation coefficients and standard error of residual deviances for each standardized predictor variable in the GLM. Estimates indicate the average change in the response variable associate with a one unit increase in the (standardized) predictor variable. Shading indicates significant association.

| Predictor | Estimate | Standard Error | P value |
| --- | --- | --- | --- |
| Intercept | -6.228 | 1.008 | <0.0005 |
| Elevation | -0.768 | 0.249 | 0.00201 |
| Bio2 | 0.983 | 0.466 | 0.0349 |
| Bio3 | 0.616 | 0.327 | 0.0600 |
| bio13 | 1.653 | 0.738 | 0.0251 |
| bio15 | -11.019 | 2.309 | <0. 0005 |
| bio19 | -4.831 | 1.107 | <0. 0005 |
| Forest Cover | 0.441 | 0.306 | 0.149 |
| VHP | - 1.256 | 0.247 | <0.0005 |

Based on the GLM, 1.4% of the study area was found to be Very Good habitat, and 11.4 % was considered Good (Table 3). Only a very small proportion of remaining habitat was within the 1 km human settlement buffer (Table 3). Suitable habitats in the southern regions of the *L. flavicauda* range run north-south through central Huánuco and Pasco, but coincide with areas where the species has been found to be absent, or where *L. l. tschudii* was found instead (this study; Aquino *et al*. 2017, 2019). In Junín, additional areas of Good habitat are present around and immediately south of the newly discovered populations. Of all Very Good and Good habitat, 17.72% and 25.95%, respectively, were found to be within Peru’s current PA network (Table 3), most of which are in PAs in the northern regions of the study area, while much of the area between the northern and Junín populations remains unprotected.

**Table 3.** Percentages of the study area that the GLM and MaxEnt model considered to be Very Good, Good and unsuitable and the amount of each habitat type within the protected area (PA) network in Peru.

| Modeling Approach | Habitat Suitability | Within Study Region (% of region- before human settlement) | Within Study Region (% of region- after human settlement) | Within PAs (% of habitat) |
| --- | --- | --- | --- | --- |
| GLM | Unsuitable | 87.10 | 87.33 | 14.32 |
|  | Good | 11.47 | 11.28 | 25.95 |
|  | Very Good | 1.43 | 1.39 | 17.72 |
| MaxEnt | Unsuitable | 86.01 | 86.2 | 13.79 |
|  | Good | 11.66 | 11.49 | 21.5 |
|  | Very Good | 2.33 | 2.31 | 38.1 |

A single iteration of the threshold-independent AUC calculation gave an AUC of 0.93 (Fig. S1). The bootstrapped evaluation gave a slightly lower mean AUC of 0.89 (95% CI 0.86-0.95). Applying the GLM to the training dataset with a 0.4 threshold produced predictions with 81.76% accuracy. The model was better able to correctly predict the pseudoabsences (86.36%) than presences (59.26%), likely due to the larger amount of pseudoabsence points in the test dataset (Table S3). The kappa statistic (0.47) indicated that the model produced predictions in good agreement with the test data, though this was not significant (p=0.70).

### MaxEnt Model

The MaxEnt model corroborated that precipitation seasonality had a strong negative correlation with species presence (Table 4). There were some differences in variable importance in comparison to the GLM results, including relatively low importance of Precipitation of Coldest Quarter (Table 2). The AUC for the MaxEnt model was 0.95 (Fig. S2) and corroborated by the mean AUC from bootstrap (0.95; 95% CI 0.93-0.97), indicating greater accuracy than the GLM.

The MaxEnt model suggested a higher percentage of Good and Very Good habitats compared to the GLM (Table 3; Fig. 4a), with percentages minimally effected after removing areas within the human settlement buffer. Suitable habitat in the southern regions of the study area included more Good habitat running north to south in Junín (Fig. 4b). Overlaying the PA network also showed more suitable habitat overall within the network compared to the GLM, with 21.5% and 38.1% of Good and Very Good being within PAs, respectively (Table 3; Fig. 5a), though much of the suitable habitat in the southern portion of the species’ range remains unprotected (Fig. 5b).

**Fig. 4.**
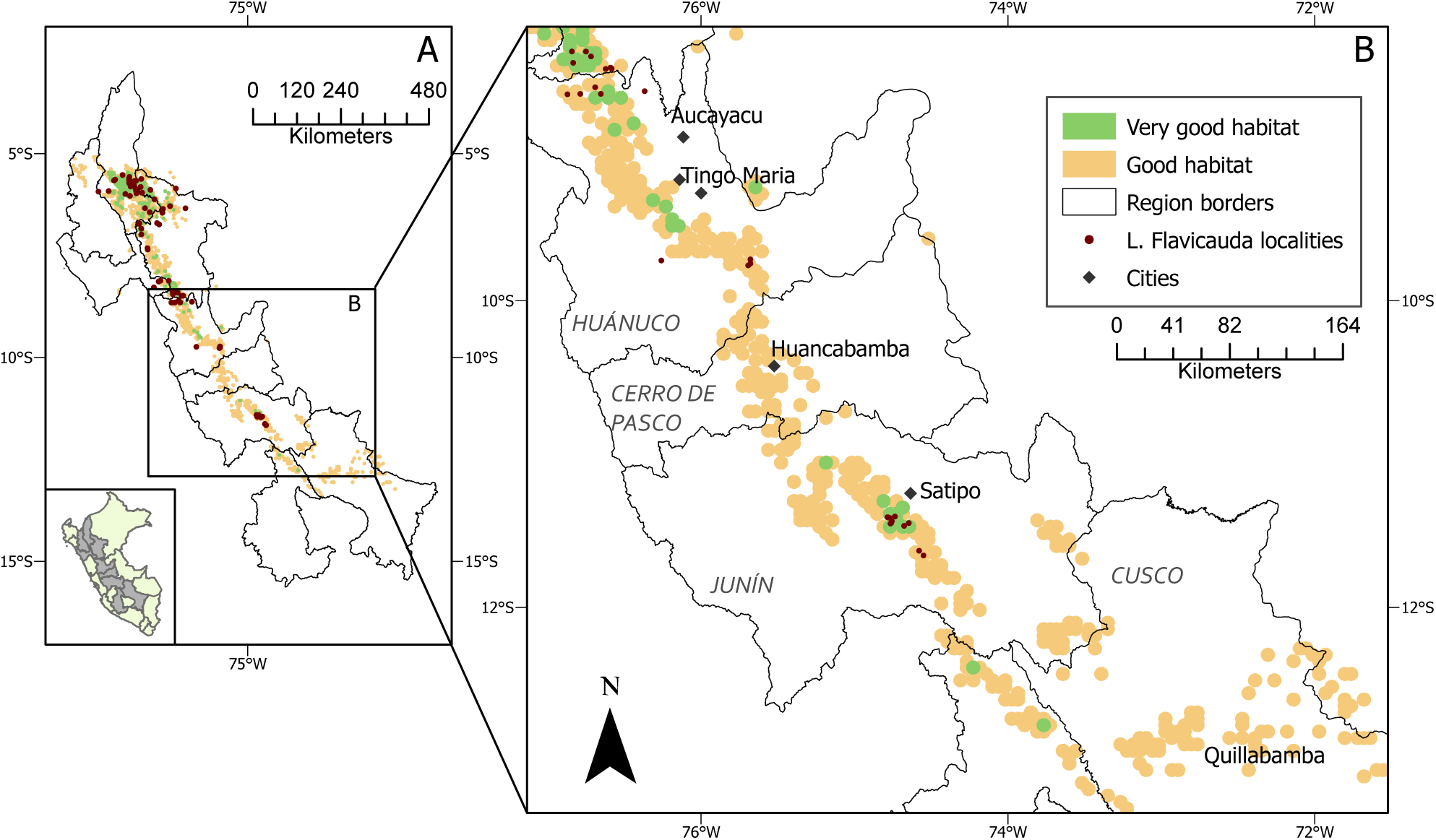
Habitat suitability maps for *L. flavicauda* as predicted by the MaxEnt model a.) throughout the described study area and b.) within the regions surrounding the southernmost *L. flavicauda* occurrences.

**Fig. 5.**
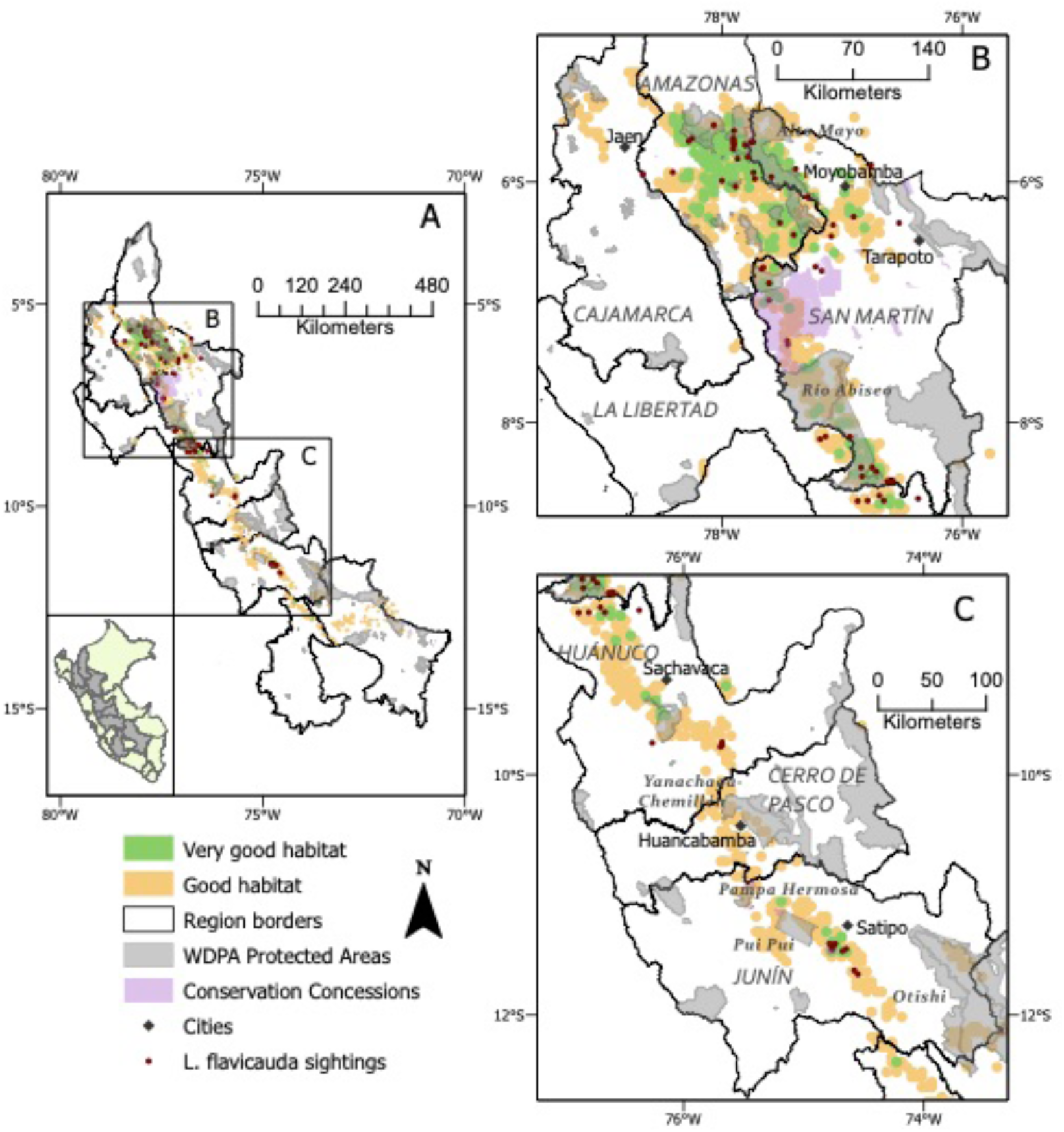
PA network coverage over habitat suitability as predicted by the Maxent model a.) over the entire study area and b.) within the regions surrounding the southernmost *L. flavicauda* occurrences

## Discussion

We carried out the first survey of *L. flavicauda* in the recently discovered southern expansion of its known range (McHugh *et al*. 2020), and our results suggest that this is a very localized population isolate of the species. Models of suitable habitat including the regions south of Huánuco and our new occurrence records provide important information on the habitat availability for the species, thereby providing insight into priority areas for focused conservation efforts and placement of conservation corridors and new protected areas. Our use of free, open-source modelling platforms will make updating and refining this analysis relatively simple with the addition of novel presence data. Our updated suitable habitats will hopefully lead to the discovery of more *L. flavicauda* populations, possibly connecting the northern and southern distributions.

Our surveys discovered multiple new sightings of *L. flavicauda* in central Junín, but only of *L. l. tschudii* in Cerro de Pasco, northern Junín and northern Cusco, and neither *Lagothrix* taxa in Ayacucho. The lack of encounters or secondary evidence of *L. flavicauda* in areas with evidence of *L. l. tschudii* may be due, in large part, to competitive exclusion. *L. l. tschudii* has been found to be one of the most abundant high-altitude species in previous surveys in Cerro de Pasco and Huánuco (Aquino *et al*. 2019), and inhabits forests with niche conditions and at elevations suitable for *L. flavicauda* (Aquino *et al*. 2016; Serrano-Villavicencio *et al*. 2021). When comparing the suitable habitat in our models with our observations of *L. l. tschudii*, we found that most are in areas well-suited to the species (Fig. 6), paralleling the work of Aquino *et al*. (2016) which indicated that *L. flavicauda* occupied similar but separate, higher-altitude habitats than *L. l. tschudii*. While competitive exclusion may explain the replacement of either *Lagothrix* taxa with its congener, further surveys are needed to rule out possible sympatry in other areas, and to confirm the absence of both taxa in Ayacucho, with future models incorporating both species.

**Fig 6.**
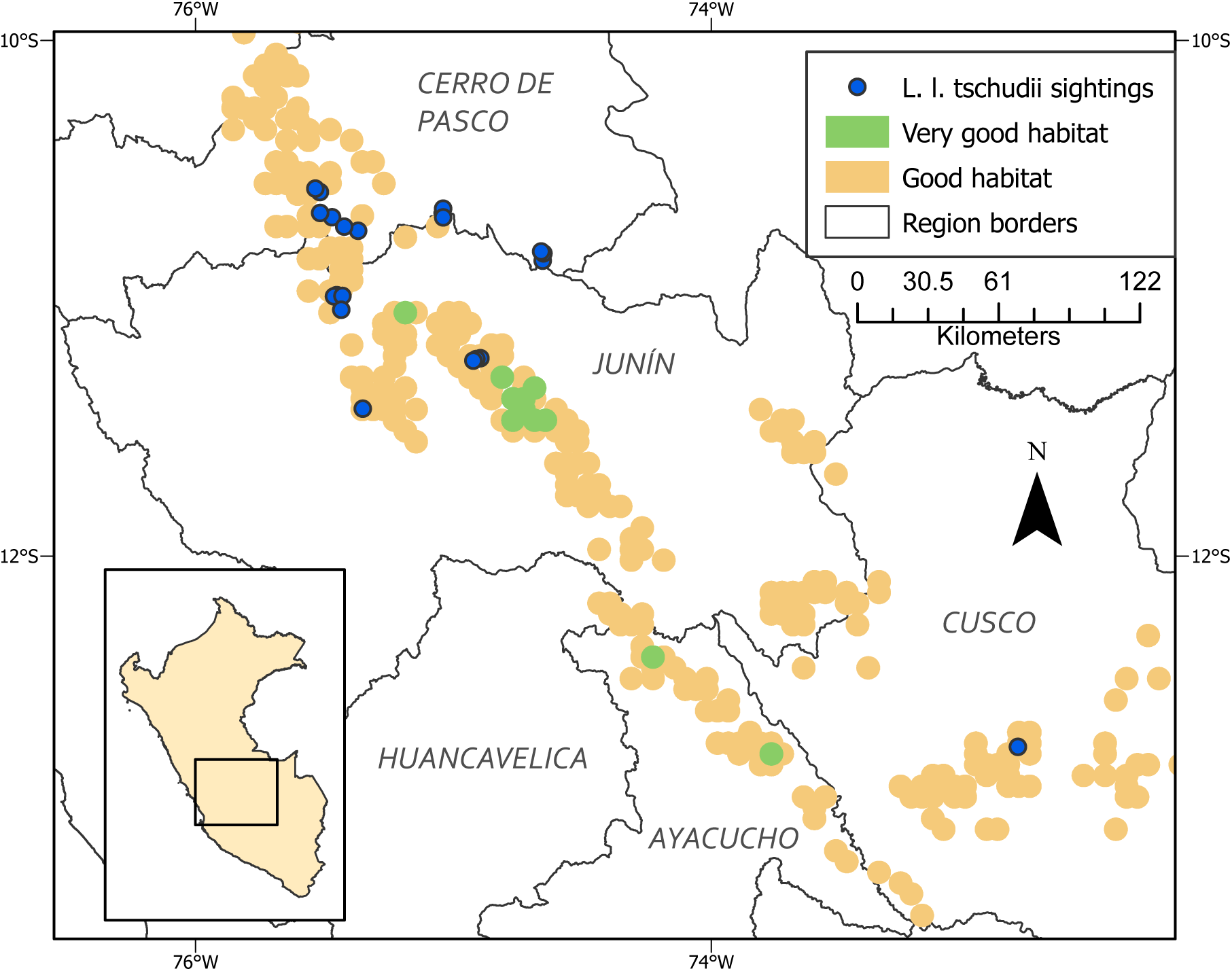
*L. l. tschudii* occurrences compared to suitable habitat predicted by the MaxEnt model. *L. flavicauda* and *L. l. tschudii* were never found occurring at the same site.

The lack of *Lagothrix* sightings in Ayacucho does not conclusively confirm absence, especially in areas where our survey efforts were limited by security issues. Almost 60% of the areas visited in this region were cultivated for coca (for cocaine production), while the remaining were small forest patches. More continuous forests did exist, but these were beyond the local jurisdictions, and controlled by armed guerrilla groups involved in drug trafficking, which we did not enter, either due to lack of permission or for our safety and that of our guides. Local informants indicated the presence of “large monkeys,” in these forests, and that they are rarely seen in the smaller patches. Informants also indicated that these monkeys were more common in previous years, but that hunting by indigenous Ashaninka groups in the area diminished their numbers. This, with further increases in hunting in combination with deforestation due to coca cultivation likely contribute to the species’ rarity. In the patches we were able to survey in these areas we only recorded the smaller bodied *Saguinus, Saimiri*, and *Cebus*. Finally, some of those interviewed in the Union Mantara and Vizcatán areas in Junín claimed, based on photographs, to have seen *L. flavicauda* in high elevation areas controlled by guerrilla groups. In particular, they pointed out the area bordered by the Río Montaro as a place where *L. flavicauda* has definitely been seen. During interviews, local people who had previously ceased coca cultivation in favour of coffee and cacao indicated a recent return to coca in the area due to low market prices for legal crops. This trend could have implications for future surveys and conservation actions.

Both of our models showed a strong negative correlation between *L. flavicauda* presence and precipitation seasonality, consistent with the results of a study by Shanee (2016) in the northern distribution of the species. This may be due to the phenology of arboreal food sources on which *L. flavicauda* depends. Though the species has been found to use a variety of food resources and plant parts, studies have shown seasonal shifts in feeding behaviour and dietary components (Shanee 2014; Shanee and Shanee 2011b). Andean species of *Cercropia*, for example, whose fruits and leaves are commonly consumed by *L. flavicauda* (Shanee 2014), experience annual reproductive changes due to climatic seasonality (Zalamea *et al*. 2011). Moreover, *L. flavicauda* have commonly been observed consuming *Ficus* spp. fruit (Leo Luna 1980; Shanee 2014; Almeyda-Zambrano *et al*. 2019), which have inter- and intra-specific asynchronous fruiting; together with abundant production, this accounts for the genus’ importance for many frugivores (Bronstein *et al*. 1990; Kattan and Valenzuela 2013). Overall, while modelling with bioclimatic variables can aid in determining areas with suitable niche conditions, these models do not provide clear explanations for a given species’ presence or absence.

Changing environmental conditions due to current and future climate change, in combination with increasing anthropogenic impacts in the TABH are predicted to decrease climatic niche availability and, in turn, species richness of plants (Ramirez-Villegas *et al*. 2014). These changes will likely lead to further habitat loss and population reductions in *L. flavicauda* and other primate species (e.g., Shanee 2016). Research suggests that the southern Peruvian Andes are experiencing a slight decrease in precipitation along with significant increases in surface temperatures (Vuille *et al*. 2003). Monthly rainfall projections based on fourth report of IPCC climate change models suggest a significant future increase in rainfall seasonality in the tropical Andes (Lavado Casimiro *et al*. 2011). Our models suggest that precipitation and seasonality are among the most important predictors of *L. flavicauda* presence, therefore such changes could have dire consequences for the species. This is especially true as lower-altitude species migrate to occupy higher elevations with climate change, increasing competition for resources (Fisher 2011). Future scenarios are further complicated by the fragmentation of habitats and intervening barriers to dispersal (Sales *et al*. 2019), which would limit *L flavicauda’s* ability to colonize new areas.

The buffer used to estimate areas as “Low” suitability based on proximity to human settlement is meant to reflect hunting pressure on *L. flavicauda* (Buckingham and Shanee 2009). Biodemographic hunting models have been created comparing human population and demographic data with primate population density to demonstrate the deleterious impact of hunting on primate “prey” species (Levi *et al*. 2011). Although hunting is known to be one of the greatest threats to *L. flavicauda* (Shanee 2012), the assumption that there is high hunting pressure closer to human settlements can be problematic due to differing practices among communities. Conversely, in areas where community conservation initiatives have been enacted, such as the community of Yambrasbamba in Amazonas, hunting of the species has been largely eliminated and populations of *L. flavicauda* have increased (Shanee *et al*. 2015). In addition to hunting, different regions in Peru have varying wildlife trafficking trends and pathways in which primates are often trafficked as pets or tourist attractions, with large-bodied primates being the most common victims (Shanee *et al*. 2017). Given this, using pressures from human hunting and trafficking activity as variables in models like this may require more detailed surveying of local practices to get a more nuanced representation of localized anthropogenic pressures.

While both models showed high statistical accuracies in their predictions, our MaxEnt model performed better. While the GLM was more conservative in the predicted amount of suitable habitat in the southern regions, there was a large amount of overlap in predictions between both models. As with all models, ours are only as accurate as the data used. We were limited by the relatively low number of localities compared to many other primate habitat suitability models (Liu *et al*. 2019). Regardless, our model predictions and survey results should encourage the continuation of surveys in southern Peru to increase presence data and model accuracy for the species.

Overlaying the PA network and the MaxEnt predictions showed that ∼75% of suitable habitat is currently unprotected, with most protection being in the northern regions of the species’ range (Fig. 5c). This is much lower than that of a gap analysis performed in 2009 (Buckingham and Shanee 2009), likely due to the recent growth of private and communal protected areas, as well as increases in state parks in the northern regions of the *L. flavicauda* range (Shanee et al. 2017; Shanee 2018; Shanee et al. 2020). Peru has 18.8% of its land covered by the current PA network, with 91.7% being within governmental PAs and 8.3% in private or communal Pas (Shanee *et al*. 2020). While national PAs generally cover larger areas, they operate at a regional scale and rarely give rights or leadership roles to local people (Shanee *et al*. 2017; Horwich *et al*. 2012). The use of protected areas is key for the conservation of *L. flavicauda*, but new state protected areas can require large areas, free of other land uses, and can take years to formally create. Private and communal PAs, on the other hand, have proven effective for conservation in the regions of Amazonas and San Martin where relatively high human population densities limit opportunities for large state-run PAs (Horwich *et al*. 2012; Shanee *et al*. 2020). In these areas, local people are the primary decision makers and enforce their own regulations, ensuring that the methods of protection are in line with the values of their community. The promotion of communally and privately managed protected areas, particularly in key corridor areas, and education across landscapes can provide much needed protection (Shanee *et al*. 2015). Further, conservation concessions cover a great amount of suitable habitat in San Martín (Fig. 5). The inclusion of the conservation concessions caused the proportion of Very Good habitat within protected areas to increase from 34.63% to 38.1 %.

In the southern portion of the species range, our analysis illustrated that only the small areas of Pampa Hermosa and Pui Pui cover an area with some suitable *L. flavicauda* habitat near the southern most populations. As such, the development of PA and community conservation initiatives in the species’ southern distribution are urgently needed, though successful conservation actions in the areas highlighted here will depend on an understanding of the socioeconomic and political situation to be able to garner local support (Chazdon *et al*. 2009).

Monitoring distributions, gene flow, and habitat suitability is critical to the conservation of *L. flavicauda*. Future research should use genetic assessments of diversity between the southern and northern populations of *L*. flavicauda to determine if gene flow is occurring. Patterns of environmental variables should also be analysed between populations to better understand factors that facilitate migration between populations, and how they could be constrained with climate change and increasing human development (Sales *et al*. 2019). Using landscape genetic methods could also determine what landscape and/or ecological features strongly facilitate or hinder gene flow between populations and be used in planning new PAs or corridors (Olah *et al*. 2016).

Further investigation of the home ranges, and resource and land use of *L. flavicauda* will allow for better interpretation of the importance of SDM predictor variables. Differences in these attributes between the northern populations and the newly-discovered Junín populations will specifically indicate the necessary factors a habitat needs to sustain viable populations. This should involve working with local communities near and within the protected areas containing suitable habitat in the south, with a particular focus on environmental education as our experience suggests a lack of understanding of the importance of animals and ecosystem function within many local immigrant (i.e., non-indigenous) communities. The presence of drug traffickers and armed groups limited which sites we could visit, particularly in the area of Vizcatán in southern Junín, thus this area remains un-surveyed, a situation which needs to be urgently rectified once conditions allow. Further successful surveys will increase the number of presence and absence points for the species, will allow for more accurate analyses of their environmental needs, and encourage the initiation of conservation efforts in these areas to continue the growth of both state and private/community led efforts for *L. flavicauda*. This will also increase the likelihood of finding other high-altitude primate species localities and help us understand their distributions and conservation needs.

## Acknowledgements

We thank the local communities we visited for allowing us to survey their surrounding forests and for their guidance in finding the monkeys. This work was funded by Boston University, and by Neotropical Primate Conservation thanks to grants from Primate Society of Great Britain, American Society of Primatologists, International Primatological Society, and Primate Conservation inc. Field work was carried out under Peruvian research permits (N° 173-2016-SERFOR-DGGSPFFS, and addendums 213-2016, 350-2017, and 389-2019). We also thank Dr. Suchi Gopal, who provided guidance in the performance of GIS analyses.

## References

Akaike, H. (1973). Information theory and extension of the Maximum Likelihood principle. Second International Symposium on Information Theory, Budapest, Hungary.

Almeyda-Zambrano, S.L. et al. (2019). Habitat preference in the critically endangered yellow tailed woolly monkey (Lagothrix flavicauda) at La Esperanza, Peru. Am. J. of Primatol. 81: e23032.

Allgas, N., Shanee, S., Peralta, A. and Shanee, N. (2014). Yellow-tailed woolly monkey (Oreonax flavicauda: Humboldt 1812) Altitudinal Range Extension, Uchiza, Perú. Neotrop. Primates 21(2): 207–208.

Aquino, R., Zárate, R., López, L., García, G. and Charpentier, E. (2015). Current status and threats to Lagothrix flavicauda and other primates in montane forest of the Región Huánuco. Primate Conserv. 29: 1–11.

Aquino, R., Garcia-Mendoza, G. and Charpentier, E. (2016). Distribution and current status of the Peruvian yellow-tailed woolly monkey (Lagothrix flavicauda) in montane forests of the Región Huánuco, Peru. Primate Conserv. 30: 31–37.

Aquino, R., García, G., Charpentier, E. and Lòpez, L. (2017). Conservation status of Lagothrix flavicauda and other primates in montane forests of San Martín and Huánuco, Peru. Rev Peru. Biol. 24(1): 025–034.

Aquino, R., López, L., Falcón, R., Díaz, S. and Gálvez, H. (2019). First inventory of primates in the montane forests of the Pasco and Ucayali regions, Peruvian Amazon. Primate Conserv. 33: 1–11

Bellamy, C., Scott, C. and Altringham, J. (2013). Multiscale, presence-only habitat suitability models: fine-resolution maps for eight bat species. J. Appl. Ecol. 50: 892–901.

Blair, M.E. and Melnick, D.J. (2012). Scale-Dependent Effects of a Heterogeneous Landscape on Genetic Differentiation in the Central American Squirrel Monkey (Saimiri oerstedii). PLoS ONE 7(8): e43027.

Bronstein J.L, Gouyon, P., Gliddon, C., Kjellberg, F. and Michaloud, G. (1990). The ecological consequences of flowering asynchrony in monoecious figs: a simulation study. Ecology 71: 2145–2156.

Butchart, S. H. M., Barnes R., Davies, C. W. N., Fernández, M. and Seddon, N. (1995) Threatened mammals of the Cordillera de Colán, Peru. Oryx 29 (4):275–281. doi:doi:10.1017/S003060530002127X

Brooks, T.M. et al. 2002. Habitat loss and extinction in the hotspots of biodiversity. Conserv. Biol. 16(4): 909–923.

Buckingham, F. and Shanee, S. (2009). Conservation priorities for the Peruvian yellow tailed woolly monkey (Oreonax flavicauda): a GIS risk assessment and gap analysis. Primate Conserv. 24: 65–71.

Case, T.J. and Taper, M.L. (2000). Interspecific competition, environmental gradients, gene flow, and the coevolution of species’ borders. Am. Nat. 155(5), 583–605.

Calkins, M.T., Beever, E.A., Boykin, K.G., Frey, J.K. and zndersen, M.C. (2012). Not-so-splendid isolation: modeling climate-mediated range collapse of a montane mammal Ochotona princeps across numerous ecoregions. Ecography 35(9), 780–791.

Chazdon, R.L., Harvey, C.A., Komar, O., Griffith, D.M., Ferguson, B.G., Martínez-Ramos, M., Morales, H., Nigh, R., Soto-Pinto, L., Van Breugal, M. and Philpott, S. M. (2009). Beyond reserves: A research agenda for conserving biodiversity in human-modified tropical landscapes. Biotropica 41(2), 142–153.

Cianfrani, C., Le Lay, G., Hirzel, A.H. and Loy, A. (2010). Do habitat suitability models reliably predict the recovery areas of threatened species? J. Appl. Ecol. 47: 421–430. 10.1111/j.1365-2664.2010.01781.x

Dai, Y., Peng, G., Wen, C., Zahoor, B., Ma, X., Hacker, C.E., and Xue, Y. (2021). Climate and land use changes shift the distribution and dispersal of two umbrella species in the Hindu Kush Himalayan region. Sci. Total Environ. 777, 146207.

Delêtre, M., Soengas, B., Jai Vidaurre, P., Isela Meneses, R., Delgado Vásquez, O., Oré Balbín, I., Santayana, M., Heider, B. and Sørensen, M. (2017). Ecotypic differentiation under farmers’ selection: molecular insight into the domestication of Pachyrhizus Rich. ex DC. (Fabaceae) in the Peruvian Andes. Evol. Appl. 10(5): 498–513.

Di Fiore, A. (2004). Diet and feeding ecology of woolly monkeys in a western Amazonian rainforest. Int. J. Primatol. 25: 767–801.

Dong, X., Chu, X., Huang, Q., Zhang, J. and Bai, W. 2019. Suitable habitat prediction Sichuan snub-nosed monlkeys (Rhinopithecus roxellana) and its implications for conservation in Baihe Nature Reserve, Sichuan, China. Environ. Sci. and Pollut. Res. 26: 32374–32384. 10.1007/s11356-019-06369-3

Dormann, C.F. et al. (2013). Collinearity: A review of methods to deal with it and a simulation study evaluating their performance. Ecography 36: 027–046.

Ewers, R.M. and Didham, R.K. (2006). Confounding factors in the detection of species responses to habitat fragmentation. Bio. Rev. 81(1), 117–142.

Hein, S., Binzenhoefer, B., Poethke, H.J., Biedermann, R., Settele, J. and Schroeder, B. (2007). The generality of habitat suitability models: A practical test with two insect groups. Basic Appl. Ecol. 8(4), 310–320.

Fack, V., Shanee, S., Vercauteren Drubbel, R., Del Viento, M., Meunier, H. and Vercauteren, M. (2018). Observation of snake (Colubridae) predation by yellow-tailed woolly monkeys (Lagothrix flavicauda) at El Toro study site, Peru. Neotrop. Primates 24 (2):79–81

Fick, S.E. and Hijmans, R.J. (2017). Worldclim 2: New 1-km spatial resolution climate surfaces for global land areas. Int. J. Climatol. 37(12): 4302–4315.

Fielding, A.H. and Bell, J.F. (1997). A Review of Methods for the Assessment of Prediction Errors in Conservation Presence/Absence Models. Environ. Conserv. 24: 38–49.

Fisher, D.O. (2011). Trajectories from extinction: where are missing mammals rediscovered? Glob. Ecol. Biogeogr. 20: 415–425.

Gallice, G.R., Larrea-Gallegos, G and Vázquez-Rowe, I. (2019). The threat of road expansion in the Peruvian Amazon. Onyx 53(2): 284–292.

GIZ (2016). Cambio de uso actual de la tierra en la Amazonía peruana: Avances e implementación en el marco de la Ley Forestal y de Fauna Silvestre 29763. Cooperación Alemana, Lima, Perú

Graves, G.R. and O’Neill, J.P (1980). Notes on the yellow-tailed woolly monkey (Lagothrix flavicauda) of Peru. J. Mammal. 61(2):345–357

Guisan, A., Thuiller, W. and Zimmermann, N.E. (2017). Habitat Suitability and Distribution Models with Applications in R. Cambridge University Press, Cambridge, UK.

Hansen, A.J., Neilson, R.P., Dale, V.H., Flather, C.H., Iverson, L.R., Currie, D.J., Shafer, S., Cook, R. and Bartlein, P.J. (2001). Global change in forests: responses of species, communities, and biomes: interactions between climate change and land use are projected to cause large shifts in biodiversity. BioScience 51(9): 765–779.

Hansen, M. C. et al. (2013). High-Resolution Global Maps of 21^st^ Century Forest Cover Change. Science 342: 850–53. Website: http://earthenginepartners.appspot.com/science-2013-global-forest. Downloaded February 2020.

Hijmans, R.J., Phillips, S., Leathwick, J. and Elith, J. (2017). R Package ‘dismo’: Species Distribution Modeling. Website: http://rspatial.org/sdm/. Accessed: February 2020.

Hijmans, R.J. et al. (2020). Package ‘raster’: Geographic data analysis and modeling. Website: https://cran.r-project.org/web/packages/raster/raster.pdf. Accessed: April 2020.

Hirzel, A.H., Le Lay, G., Helfer, V., Randin, C., and Guisan, A. (2006). Evaluating the ability of habitat suitability models to predict species presences. Ecol. Modell. 199(2), 142–152.

Horwich, R.H., Lyon, J., Bose, A. and Jones, C.B. (2012). Preserving biodiversity and ecosystems: catalyzing contagion. In Deforestation Around the World: 283–318. Moutinho, P. (Ed.). Rijeka: InTech.

Humanitarian OpenStreetMap Team. (2020). United Nations Office for the Coordination of Humanitarian Affairs. Website: https://data.humdata.org/dataset/hotosm_per_buildings. Downloaded February 20, 2020.

Hurvich, C.M. and Tsai, C. (1993). A corrected Akaike information criterion for vector autoregressive model selection. J. Time Ser. Anal. 14(3): 271–279.

Kattan, G.H. and Valenzuela, L.A. (2013). Phenology, abundance and consumers of figs (Ficus spp.) in a tropical cloud forest: evaluation of a potential keystone resource. J. Trop. Ecol. 29(5): 401–407.

Lagothrix flavicauda (Humboldt, 1812) in GBIF Secretariat (2021). GBIF Backbone Taxonomy. Checklist dataset https://doi.org/10.15468/dl.xmcds9 accessed via http://GBIF.org on 2021-09-04.

LaRue, M.A. and Nielsen, C.K. 2008. Modelling potential dispersal corridors for cougars in midwestern North America using least-cost path methods. Ecol. Modell. 212: 372–381.

Laurance, W. F. (2018). Conservation and the global infrastructure tsunami: Disclose, debate, delay!. Trends Ecol. Evol. 33(8), 568–571.

Lavado Casimiro, W.S., Labat, D., Guyot, J.L. and Ardoin-Bardin, S. (2011). Assessment of climate change impacts on the hydrology of the Peruvian Amazon-Andes basin. Hydrol. Process. 25: 3721–3734.

Leo Luna, M. (1980). First field study of the yellow-tailed woolly monkey. Oryx 15: 386–389.

Levi, T., Shepard, G.H., Ohl-Schacherer, J., Wilmer, C. C., Peres, C. A. and Yu, D. W. (2011). Spatial tools for modeling the sustainability of subsistence hunting in tropical forests. Ecol. Appl. 21(5): 1802–1818.

Liu, J., Fitzgerald, M., Liao, H., Luo, Y., Jin, Y., Li, X., Yang, X., Hirata, S. and Matsuzawa, T. (2019). Modeling habitat suitability for Yunnan snub-nosed monkeys in Laojun Mountain National Park. Primates 10.1007/s10329-01900776-3

McHugh, S.M., F.M. Cornejo, J. McKibben, M. Zarate, T. Carlos, C.F. Jiménez and C.A. Schmitt. (2020). First detection and updated geographic range of the Peruvian yellow-tailed woolly monkey (Lagothrix flavicauda) in the region Junín, Peru. Oryx 54(6): 814–818. DOI: 0.1017/S003060531900084X.

Mittermeier R.A, de Macedo-Ruiz, H, Luscombe, A (1975). A woolly monkey rediscovered in Peru. Oryx 13 (1):41–46

Newbold, T. et al. (2014). A global model of the response of tropical and sub-tropical forest biodiversity to anthropogenic pressures. Proc. R. Soc. B: Biol. Sci. 281(1792), 20141371

NOAA (National Oceanic and Atmospheric Administration). 2020. Climate Prediction Center. Website:https://www.cpc.ncep.noaa.gov/products/assessments/assess_97/samer.html. Accessed: June 2020.

Olah, G., Smith, A.L., Asner, G.P., Brightsmith, D.J., Heinsohn, R.G. and Peakall, R. (2017). Exploring dispersal barriers using landscape genetic resistance modelling in scarlet macaws of the Peruvian Amazon. Landsc. Ecol. 32: 445–456.

Oliveira, P.J.C., Asner, G.P., Knapp, D.E., Almeyda, A., Galván-Gildemeister, R., Keene, S., Raybin, R.F., Smith, R.C. (2007). Land-Use Allocation Protects the Peruvian Amazon. Science 317 (5842):1233. doi:10.1126/science.1146324

Parker, T.A and Barkley, L.J (1981). New locality for the yellow-tailed woolly monkey. Oryx 16 (1):71–71

Pásztor, L., Barabás, G. and Meszéna, G. (2020). Competitive Exclusion and Evolution: Convergence Almost Never Produces Ecologically Equivalent Species: (A Comment on McPeek,“Limiting Similarity? The Ecological Dynamics of Natural Selection among Resources and Consumers Caused by Both Apparent and Resource Competition”). Am. Nat. 195(4), E112–E117.

Patterson, B.D. and López Wong, C. (2014) Mamíferos/Mammals. In Peru: Cordillera Escalera. Rapid Biological Inventories Report 26: 154–167. Pitman, N et al. (Ed.). Chicago: The Field Museum.

Phillips, S., Anderson, R.P. and Schapire, R.E. (2006). Maximum entropy modeling of species geographic distributions. Ecol. Modell. 190(3-4): 231–259.

Phillips, S.J., Dudík, M., Elith, J., Graham, C. H., Lehmann, A., Leathwick, J., Ferrier, S. (2009). Sample selection bias and presence-only distribution models: implications for background and pseudo-absence data. Ecol. Appl. 19: 181–197.

Programa Bosques (2015) Bosque - no bosque y pérdida de bosque Húmedo Amazónico 2000-2014. Programa Nacional de Conservación de Bosques para la Mitigación del Cambio Climatico, Lima, Peru.

R Core Team. (2018). R: A language and environment for statistical computing. R Foundation for Statistical Computing, Vienna, Austria. Website: https://www.Rproject.org/.

Ramirez-Villegas, J., Cuesta, F., Devenish, C., Peralvo, M., Jarvis, A. and Arnillas, C.A. (2014). Using species distribution models for designing conservation strategies of Tropical Andean biodiversity under climate change. J. Nat. Conserv. 22: 391–404.

Robillard, C.M., Coristine, L.E., Soares, R.N. and Kerr, J.T. (2015). Facilitating climate-change-induced range shifts across continental land-use barriers. Conserv. Biol. 29(6), 1586–1595.

Rodriguez, E., Morris, C.S., Belz, J.E., Chapin, E.C., Martin, J.M., Daffer, W., and Hensley, S. (2005). An assessment of the SRTM topographic products, Technical Report JPL D-31639. Jet Propulsion Laboratory, pp 143. Pasadena, California. Website: http://srtm.csi.cgiar.org/srtmdata/. Downloaded February 2020.

Sales, L.P., Ribeiro, B.R., Pires, M.M., Chapman, C.A. and Loyola, R. (2019). Recalculating route: dispersal constraints will drive the redistribution of Amazon primates in the Anthropocene. Ecography 42: 1789–1801.

Serrano-Villavicencio, J.E., Shanee, S. and Pacheco, V. (2021). Lagothrix flavicauda (Primates: Atelidae). Mamm. Species 53(1010) 134–144. DOI: 10.1093/mspecies/seab013

Shanee, N., Shanee, S. and Maldonado, A.M. (2007). Conservation assessment and planning for the yellow-tailed woolly monkey (Oreonax flavicauda) in Peru. Wildlife Biol. 3: 73-82.

Shanee, S. and Shanee, N. (2009). A new conservation NGO, neotropical primate conservation: project experiences in Perú. Int. NGO J. 4(7): 329–332.

Shanee, S. (2011). Distribution survey and threat assessment of the yellow-tailed woolly monkey (Oreonax flavicauda; Humboldt 1812), northeastern Peru. Int. J. Primatol. 32: 691–707.

Shanee, S. and Shanee N. (2011a). Population density estimates for the critically endangered yellow-tailed woolly monkey (Lagothrix flavicauda) at La Esperanza, northeastern Peru. Int. J. Primatol. 32: 691–180.

Shanee, S. and Shanee, N. (2011b). Activity budget and behavioural patterns of free-ranging yellow-tailed woolly monkeys Oreonax flavicauda (Mammalia: Primates), at La Esperanza, northeastern Peru. Contrib. Zool. 80:269–277.

Shanee, N. (2012) Trends in local wildlife hunting, trade and control in the Tropical Andes Hotspot, Northeastern Peru. Endangered Species Research 19 (2):177–186

Shanee, S. (2014). Ranging behaviour, daily path lengths, diet and habitat use of yellow tailed woolly monkeys (Oreonax flavicauda) at La Esperanza, Peru. In The woolly monkey: behaviour, ecology, conservation and systematics: 167–186. Defler, T. R and P. R. Stevenson (Ed.). New York: Springer Verlag.

Shanee, N., Shanee, S. and Horwich, R.H. (2015). Effectiveness of locally run conservation initiatives in north-east Peru. Oryx 49(2): 239–247.

Shanee, S. and Shanee, N. (2015). Measuring success in a community conservation project: local population increase in a critically endangered primate, the yellow-tailed woolly monkey (Lagothrix flavicauda) at La Esperanza, northeastern Peru. Trop. Conserv. Sci. 8(1): 169–186.

Shanee, S. (2016). Predicting Future Effects of Multiple Drivers of Extinction Risk in Peru’s Endemic Primate Fauna. In Ethnoprimatology, Developments in Primatology, Progress and Prospects: 315–349. Waller, M. T. (Ed.), Switzerland: Springer International Publishing.

Shanee, N. and Shanee, S. (2016). Land trafficking, migration and conservation in the “No Man’s Land” of northeastern Peru. Trop. Conserv. Sci. 9(4): 1–16.

Shanee, N., Mendoza, A.P. and Shanee, S. (2017). Diagnostic overview of the illegal trade in primates and law enforcement in Peru. Am. J. Primatol. 79 (11): e22516. doi:10.1002/ajp.22516

Shanee, S., Shanee, N., Monteferri, B., Allgas, N., Pardo, A.A. and Horwich, R.H. (2017). Protected area coverage of threatened vertebrates and ecoregions in Peru: Comparison of communal, private and state reserves. J. Environ. Manage. 202, 12–20.

Shanee, S. Allgas, N. and Shanee, N. (2018). Community conservation as a tool for primate conservation in Peru. In Primatology, Biocultural Diversity and Sustainable Development in Tropical Forests: 320–329. Rommens, D. and J. Pulido Mata (Ed.). Mexico: UNESCO.

Shanee, S., Shanee, N., Lock, W. and Espejo-Uribe, M.J. (2020). The development and growth of non-governmental conservation in Peru: Privately and communally protected areas. Hum. Ecol. 48: 681–693.

Shanee, S., Cornejo, F.M., Aquino, R., Mittermeier, R.A. & Vermeer, J. (2021). Lagothrix flavicauda (amended version of 2019 assessment). The IUCN Red List of Threatened Species 2021: e.T39924A192307818. https://dx.doi.org/10.2305/IUCN.UK.2021-1.RLTS.T39924A192307818.en. Accessed on 31 January 2022.

Solórzano-García, B., Zubillaga, D., Piñero, D. and Vázquez-Domínguez, E. (2021). Conservation implications of living in forest remnants: inbreeding and genetic structure of the northernmost mantled howler monkeys. Biotropica 53(4), 1163–1177.

Strier, K.N. (1992). Atelinae adaptations: behavioral strategies and ecological constraints. Am. J. Phys. Anthropol. 88: 515–524.

UNEP-WCMC. 2020. Protected Area Profile for Peru from the World Database of Protected Areas, June 2020. Available at: http://www.protectedplanet.net

Vermote, E. (2019). NOAA CDR Program. NOAA Climate Data Record (CDR) of AVHRR Normalized Difference Vegetation Index (NDVI), Version 5. Vegetation Health Product. NOAA National Centers for Environmental Information. https://doi.org/10.7289/V5ZG6QH9. Downloaded May 20, 2020.

Vuille, M., Bradley, R.S., Werner, M. and Keimig, F. (2003). 20th century climate change in the Tropical Andes: observations and model results. In Variability and Change in High Elevation Regions: Past, Present & Future. Advances in Global Change Research, vol 15: 75–99. Diaz, H. F. (Ed.). Dordrecht: Springer.

Xiao, H., McDonald-Madden, E., Sabbadin, R., Peyrard, N., Dee, L. E. and Chadès, I. (2019). The value of understanding feedbacks from ecosystem functions to species for managing ecosystems. Nat. Comm. 10(1), 1–10.

Young, K.R. (1996). Threats to biological diversity caused by coca/cocaine deforestation in Peru. Environ. Conserv. 23 (1):7–15. doi:doi:10.1017/S0376892900038200

Zalamea, P., Munoz, F., Stevenson, P.R., Paine, C.E.T., Sarmiento, C., Sabatier, D. and Heuret, P. (2011). Continental-scale patterns of Cecropia reproductive phenology: evidence from herbarium specimens. Proceedings of the Royal Society B 278: 2437–2445.

