## Supplemental Materials for "Expanded Distribution and Predicted Suitable Habitat for the Critically Endangered Yellow-tailed Woolly Monkey (*Lagothrix flavicauda*) in Peru"

**Table S1.** List of presence data used in the creation of habitat suitability models (HSMs) coming from published literature, Global Biodiversity Information Facility, and the surveys in this study.

| **Longitude** | **Latitude** | **Source** |
| --- | --- | --- |
| -78.658683 | -5.9337166 | Shanee 2011 |
| -77.1706 | -6.7379334 | Shanee 2011 |
| -77.2216 | -6.7047333 | Shanee 2011 |
| -76.904117 | -6.28905 | Shanee 2011 |
| -76.5264 | -6.3403167 | Shanee 2011 |
| -76.259567 | -9.7376667 | Shanee 2011 |
| -77.670083 | -6.7216166 | Shanee 2011 |
| -76.622117 | -8.4885833 | Shanee 2011 |
| -76.84255 | -8.37455 | Shanee 2011 |
| -77.608633 | -6.8414167 | Shanee 2011 |
| -77.654833 | -6.7099334 | Shanee 2011 |
| -77.403283 | -6.4377833 | Shanee 2011 |
| -78.4075 | -5.9181167 | Shanee 2011 |
| -78.067367 | -5.5250667 | Shanee 2011 |
| -77.5182 | -6.3391167 | Shanee 2011 |
| -78.277 | -5.6603002 | Shanee 2011 |
| -77.78545 | -5.7906 | Shanee 2011 |
| -78.00025 | -5.9896834 | Shanee 2011 |
| -77.870583 | -5.8066167 | Shanee 2011 |
| -77.726333 | -5.9733833 | Shanee 2011 |
| -78.658683 | -5.9337166 | Shanee 2011 |
| -77.8863 | -6.0369167 | Shanee 2011 |
| -77.806433 | -5.69205 | Shanee 2011 |
| -78.2506 | -5.6383333 | Shanee 2011 |
| -77.9 | -5.5720667 | Shanee 2011 |
| -77.90535 | -5.6356167 | Shanee 2011 |
| -77.901267 | -5.65915 | Shanee 2011 |
| -77.906 | -5.6594833 | Shanee 2011 |
| -77.900017 | -5.68785 | Shanee 2011 |
| -77.903717 | -5.6868167 | Shanee 2011 |
| -77.9076 | -5.7236167 | Shanee 2011 |
| -77.7373 | -5.9210166 | Shanee 2011 |
| -77.456583 | -7.31905 | Shanee 2011 |
| -77.385867 | -5.89015 | Shanee 2011 |
| -77.28705 | -6.1214167 | Shanee 2011 |
| -77.07535 | -6.357 | Shanee 2011 |
| -77.09105 | -6.4511167 | Shanee 2011 |
| -77.452483 | -7.3509167 | Shanee 2011 |
| -76.7508 | -8.3751167 | Shanee 2011 |
| -76.71785 | -8.4067833 | Shanee 2011 |
| -77.758717 | -5.66975 | Shanee 2011 |
| -77.74125 | -5.6674667 | Shanee 2011 |
| -77.589683 | -5.95725 | Shanee 2011 |
| -76.87165 | -8.6538333 | Shanee 2011 |
| -76.788 | -8.6510667 | Shanee 2011 |
| -76.690933 | -8.608 | Shanee 2011 |
| -77.138033 | -8.1170167 | Shanee 2011 |
| -77.184167 | -8.1384666 | Shanee 2011 |
| -74.785461 | -11.411369 | McHugh *et al* 2019 |
| -74.784663 | -11.412653 | McHugh *et al* 2019 |
| -74.773875 | -11.414683 | McHugh *et al* 2019 |
| -74.77765 | -11.41355 | McHugh *et al* 2019 |
| -76.59025 | -8.4797333 | Allgas *et al* 2014 |
| -76.760528 | -5.8561111 | Allgas *et al* 2014 |
| -75.693489 | -9.7696277 | Aquino *et al* 2017 |
| -76.368088 | -8.6336612 | Aquino *et al* 2017 |
| -76.654028 | -8.6509522 | Aquino *et al* 2017 |
| -76.583391 | -8.4858371 | Aquino *et al* 2017 |
| -76.834178 | -8.4486378 | Aquino *et al* 2017 |
| -76.935172 | -8.1195675 | Aquino *et al* 2017 |
| -77.608898 | -6.9833265 | Aquino *et al* 2017 |
| -75.678877 | -9.7577732 | Aquino *et al* 2016 |
| -75.678958 | -9.7299714 | Aquino *et al* 2016 |
| -74.765884 | -11.453018 | This study |
| -74.756811 | -11.437754 | This study |
| -74.760899 | -11.448999 | This study |
| -74.646722 | -11.449616 | This study |
| -74.677038 | -11.46781 | This study |
| -74.760899 | -11.448999 | This study |
| -74.550389 | -11.660964 | This study |
| -74.579308 | -11.630088 | This study |
| -74.737175 | -11.406908 | This study |
| -77.746516 | -5.862236 | GBIF |
| -77.624564 | -5.818765 | GBIF |
| -77.907123 | -5.59823 | GBIF |
| -77.610061 | -5.623244 | GBIF |
| -77.3 | -8.28 | GBIF |
| -77.77 | -5.67 | GBIF |

**Table S2** Data layers used to as predictor variables to build the species distribution models for *L. flavicauda*. To apply the model to the study region and generation predictions of species presence probability, all layers with the exception of the buildings layer were aggregated to the smallest resolution (30 meters).

| Variable Category | Variable Name | Description | Original Data Type | Source |
| --- | --- | --- | --- | --- |
| Climate | bio1 | Annual Mean Temperature | Raster (1 kilometer resolution) | WorldClim Global Climate Data (Fick and Hijmans 2017) |
|  | bio2 | Mean Diurnal Range (Mean of monthly (max temp - min temp)) |  |  |
|  | bio3 | Isothermality (BIO2/BIO7) (* 100) |  |  |
|  | bio4 | Temperature Seasonality (standard deviation *100) |  |  |
|  | bio5 | Max Temperature of Warmest Month |  |  |
|  | bio6 | Min Temperature of Coldest Month |  |  |
|  | bio7 | Temperature Annual Range (BIO5-BIO6) |  |  |
|  | bio8 | Mean Temperature of Wettest Quarter |  |  |
|  | bio9 | Mean Temperature of Driest Quarter |  |  |
|  | bio10 | Mean Temperature of Warmest Quarter |  |  |
|  | bio11 | Mean Temperature of Coldest Quarter |  |  |
|  | bio12 | Annual Precipitation |  |  |
|  | bio13 | Precipitation of Wettest Month |  |  |
|  | bio14 | Precipitation of Driest Month |  |  |
|  | bio15 | Precipitation Seasonality (Coefficient of Variation) |  |  |
|  | bio16 | Precipitation of Wettest Quarter |  |  |
|  | bio17 | Precipitation of Driest Quarter |  |  |
|  | bio18 | Precipitation of Warmest Quarter |  |  |
|  | bio19 | Precipitation of Coldest Quarter |  |  |
| Elevation | Elevation | Digital elevation data (in meters) | Raster (30 meter resolution) | NASA Shuttle Radar Topography Mission (Rodriguez et al. 2005) |
| Forest | 2000 Forest Cover | Percent forest cover as a result of Landsat 8 OLI data in the year 2000 | Raster (30 meter resolution) | Global Forest Watch (Hansen et al. 2013) |
|  | Forest Loss 2000-2019 | Year of loss/no forest cover loss between 2000-2019 as a result of Landsat 8 OLI data in the year 2000 |  |  |
|  | Forest gain 2000-2012 | Binary outcome of gain/no gain of forest cover between 2000-2012 as a result of Landsat 8 OLI data in the year 2000 |  |  |
| Vegetation | Blended Vegetation Health Product | A proxy for vegetation health calculated by Advanced Very High Resolution Radiometer (AVHRR) products and the Global Area Coverag (GAC) dataset. | Raster (1 kilometer resolution) | NOAA Centre for Satellite Applications and Research (Vermote 2019) |
| Anthropogenic | Distance from Urban Areas | Distance from buffers around buildings from Humanitarian Open Street Map Team | Vector | United Nations Office for the Coordination of Humanitarian Affairs (Humanitarian OpenStreetMap Team 2020) |
| Protected Areas | Protected Area Network in Peru | All nationally reported regional, national, private, and communal reserves and protected areas in Peru. | Vector | World Database of Protected Areas (UNEP-WCMC 2020) |
|  | Conservation Concessions in Peru | All reported conservation concession areas in Peru. | Vector | Peruvian National Forestry Service (SERFOR) |

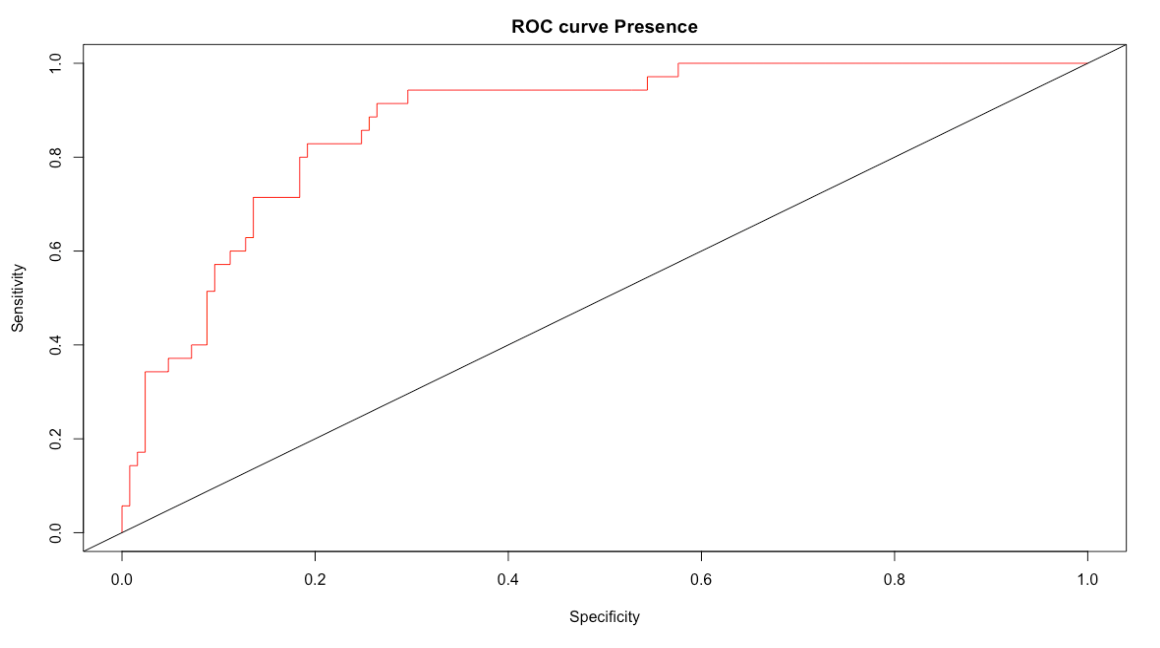

**Figure S1.** The receiving operator characteristic (ROC) curve produced by the GLM predictions against a 40% test dataset of *L. flavicauda* presence and pseudoabsences. The ROC (red; AUC = 0.930) is compared to the diagonal representing random guess (black; AUC = 0.5). Sensitivity represents accurate predicted presence probability while specificity represents accurate true absence probability.

**Table S3.** Confusion matrix of the predicted presence and pseudoabsence points produced by HSM25 against actual recorded points with a 0.4 presence probability threshold of predictions. Predictions were calculated by building the model with 60% of the data (training set) and applying the model to a 40% test dataset of *L. flavicauda* presences and pseudoabsence.

|  | Recorded | | |  |
| --- | --- | --- | --- | --- |
| Predicted |  | Pseudoabsence | Presence | Totals |
|  | Pseudoabsence | 114 | 11 | 125 |
|  | Presence | 18 | 16 | 34 |
|  | Totals | 132 | 27 | 159 |

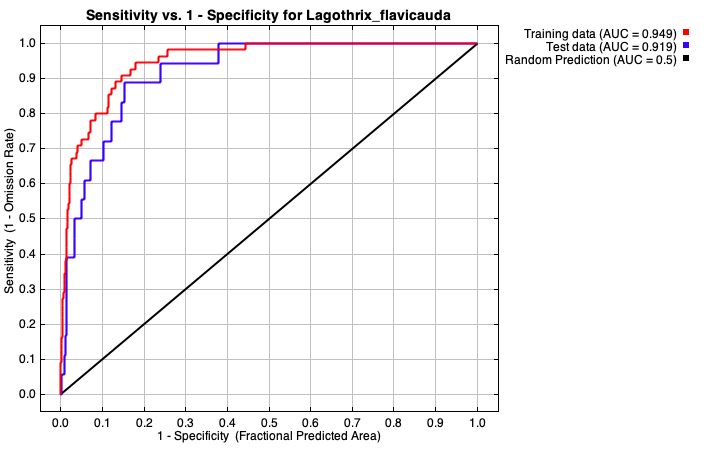

**Figure S2.** ROC curve as calculated by MaxEnt program software. Repeating the calculation corroborated the curve and AUC value.
